# High glucose treatment induced nuclei aggregation of microvascular endothelial cells via foxo1a-klf2a pathway

**DOI:** 10.1101/2024.04.29.591787

**Authors:** Xiaoning Wang, Xinyi Kang, Bowen Li, Changshen Chen, Liping Chen, Dong Liu

## Abstract

**BACKGROUND:** Hyperglycemia is a major contributor to endothelial dysfunction and blood vessel damage, leading to severe diabetic microvascular complications. Despite the growing body of research on the underlying mechanisms of endothelial cell dysfunction, the available drugs based on current knowledge fall short of effectively alleviating these complications. Therefore, our endeavor to explore novel insights into the cellular and molecular mechanisms of endothelial dysfunction is crucial for the field.

**METHODS:** In this study, we carried out a high-resolution imaging and time-lapse imaging analysis of the behavior of endothelial cells *in Tg(kdrl:ras-mCherry::fli1a:nEGFP)* zebrafish embryos upon high glucose treatment. Genetic manipulation and chemical biology approaches were utilized to analyze the underlying mechanism of high-glucose-induced nuclei aggregation and aberrant migration of zebrafish endothelial cells and cultured human endothelial cells. Bioinformatical analysis of single-cell RNA sequencing data and molecular biological techniques to identify the target genes of Foxo1a.

**RESULTS:** In this study, we observed that the high glucose treatment resulted in nuclei aggregation of endothelial cells in zebrafish intersegmental vessels (ISVs). Additionally, the aberrant migration of microvascular endothelial cells in high glucose-treated embryos, which might be a cause of nuclei aggregation, was discovered. High glucose-induced aggregation of vascular endothelial nuclei via foxo1a downregulation in zebrafish embryos. Then, we revealed that high glucose resulted in the downregulation of foxo1a expression and increased the expression of its direct downstream effector, klf2a, through which the aberrant migration and aggregation of vascular endothelial nuclei were caused.

**CONCLUSIONS:** High glucose treatment caused the nuclei of endothelial cells to aggregate *in vivo*, which resembles the crowded nuclei of endothelial cells in microaneurysms. High glucose suppresses foxo1a expression and increases the expression of its downstream effector, klf2a, thereby causing the aberrant migration and aggregation of vascular endothelial nuclei. Our findings provide a novel insight into the mechanism of microvascular complications in hyperglycemia.

## Introduction

Diabetic microvascular complications include diabetic retinopathy, diabetic nephropathy, and diabetic neuropathy [1], which might be potentially caused by tissue exposure to chronic hyperglycemia. High blood glucose levels harm endothelial cells (EC), leading to endothelial “dysfunction,” resulting in microvascular hyperplasia, vascular lumen narrowing, and other pathological manifestations. The molecular mechanism of endothelial dysfunction in diabetes is complex, involving multiple factors, such as oxidative stress [2–5], activation of PKC [6, 7], overexpression of growth factors [8–10], nonenzymatic glycation of proteins [11, 12], impaired insulin activation of PIP-3 kinase [13–15], and others. While cumulating studies have shed light on the underlying mechanisms of endothelial cell dysfunction, there remains a significant knowledge gap regarding the causes and mechanisms of diabetic microvascular complications. Notably, currently available drugs cannot relieve these vascular diseases satisfactorily [16–21]. Therefore, further exploration of the cellular and molecular drivers of endothelial dysfunction is crucial for developing effective therapeutic strategies for diabetic microvascular disease.

Zebrafish is an advantageous model and has been widely used in vascular research. Zebrafish embryos are optically transparent, allowing high-resolution live imaging of blood vessel development and pathological processes [22]. Due to its similar glucose metabolism pathways to humans, the zebrafish is also an emerging disease model organism for research on diabetes and its vascular complications [23]. Moreover, the genetic manipulation strategies of zebrafish are relatively simple, making it very convenient for gene loss of function and gain of function [24–27]. The *in vivo* imaging analysis of the zebrafish model offers an opportunity to discover novel behaviors of microvascular endothelial cells under hyperglycemia conditions.

In the present study, we show that high glucose treatment induces the aggregation of vascular endothelial nuclei in zebrafish embryos’ ISVs. Furthermore, we investigate the molecular mechanisms under the phenotypes. This study showed for the first time that high glucose causes the aggregation of endothelial nuclei, providing a novel insight into the mechanism of microvascular complications in hyperglycemia.

## Results

### High glucose treatment induced the aggregation of vascular endothelial nuclei in the ISVs of zebrafish embryos

In the previous work, we reported that high glucose treatment could cause Intersegmental vessel (ISV) hyperplasia in zebrafish embryos [28]. To further examine the effects of high glucose on endothelial cells, we treated zebrafish embryos with a high dose of D-glucose (300 mM) for a short period. We subsequently measured the glucose concentration in the embryos, and the results showed that the glucose concentration in the embryos treated with high glucose was significantly higher than that in the control group (Supplementary Figure 1). Confocal imaging analysis of *Tg(kdrl:ras-mCherry::fli1a:nEGFP)* embryos revealed that the treatment of high glucose from 48 to 72 hpf caused endothelial nucleus aggregation in almost all ISVs of the embryos (Figure 1a-h). The nearest neighbor distance (NND) of ECs nuclei and distance from the ECs nuclei to the midline (DNTM) were also reduced compared with control (Figure 1i, k). Subsequently, the ISVs were partitioned into four longitudinal zones, wherein the EC nuclei in the control group exhibited uniform distribution (Figure 1e, j). In contrast, those in the treated group were predominantly concentrated in B and C zones (Figure 1h, j). The range and standard deviation (SD) of the DNTM in the treated embryos were also smaller than those in the control siblings (Figure 1l-m). Furthermore, we conducted the treatments on the embryos during various time intervals and observed that exposure to high glucose for either 1 day (24∼48 hpf, 48∼72 hpf, 72∼96 hpf) or 2 days (48∼96 hpf), all led to the aggregation of endothelial nuclei and decreased NND in the ISVs of zebrafish embryos (Figure 2a-d) in comparison to control (Figure 2a’-d’). These results suggest that the endothelial nuclei in ISVs become more concentrated after high glucose treatment.

**Figure 1.**
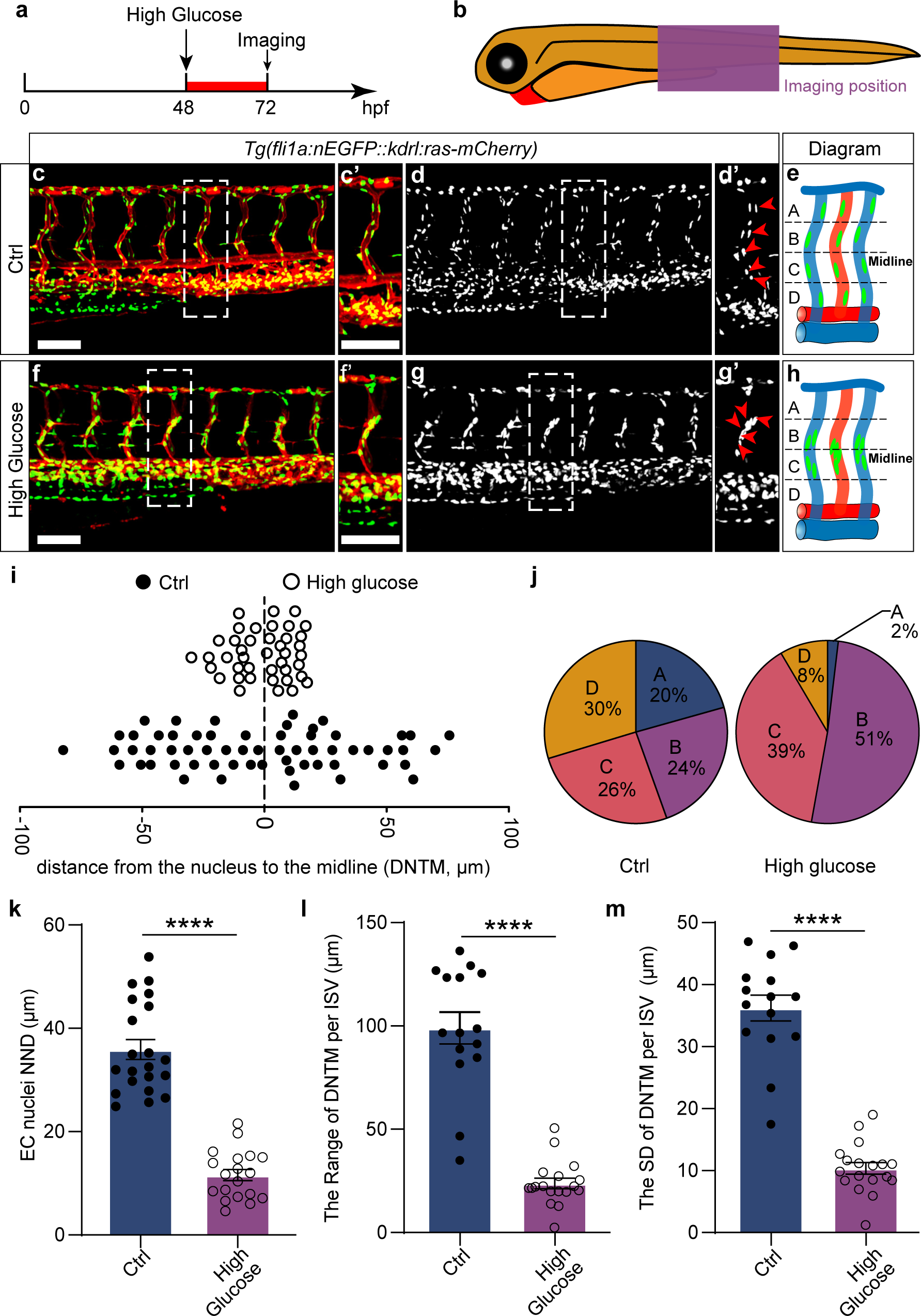
High glucose shock treatment induced the aggregation of vascular endothelial cell nuclei in the ISVs of zebrafish embryos. (a,b) Schematic diagram showing the high glucose shock treatment timeline and confocal imaging region. (c-g) Confocal imaging analysis of control and high glucose shock treated *Tg(fli1a:nEGFP::kdrl:ras-mCherry)* embryos at 72 hpf. (c’-g’) The magnifications of the red dotted boxes in c-g, respectively. The arrowheads indicate the endothelial cell nuclei in the ISV. (i) The distance from the ECs nuclei to the midline (DNTM) in ISVs in control and high glucose shock treated embryos. (j) The proportion of ECs nuclei in ISVs in the four zones is shown in panels e and h. (k) Statistics of the nearest neighbor distance (NND) of ECs nuclei in control embryos and high glucose treated embryos. *t*-test. (i,m) Statistics of the range and standard deviation (SD) of DNTM in control embryos and high glucose-treated embryos. *t*-test. \*\*\*\**p*<0.0001. Scale bars, 100 µm.

**Figure 2.**
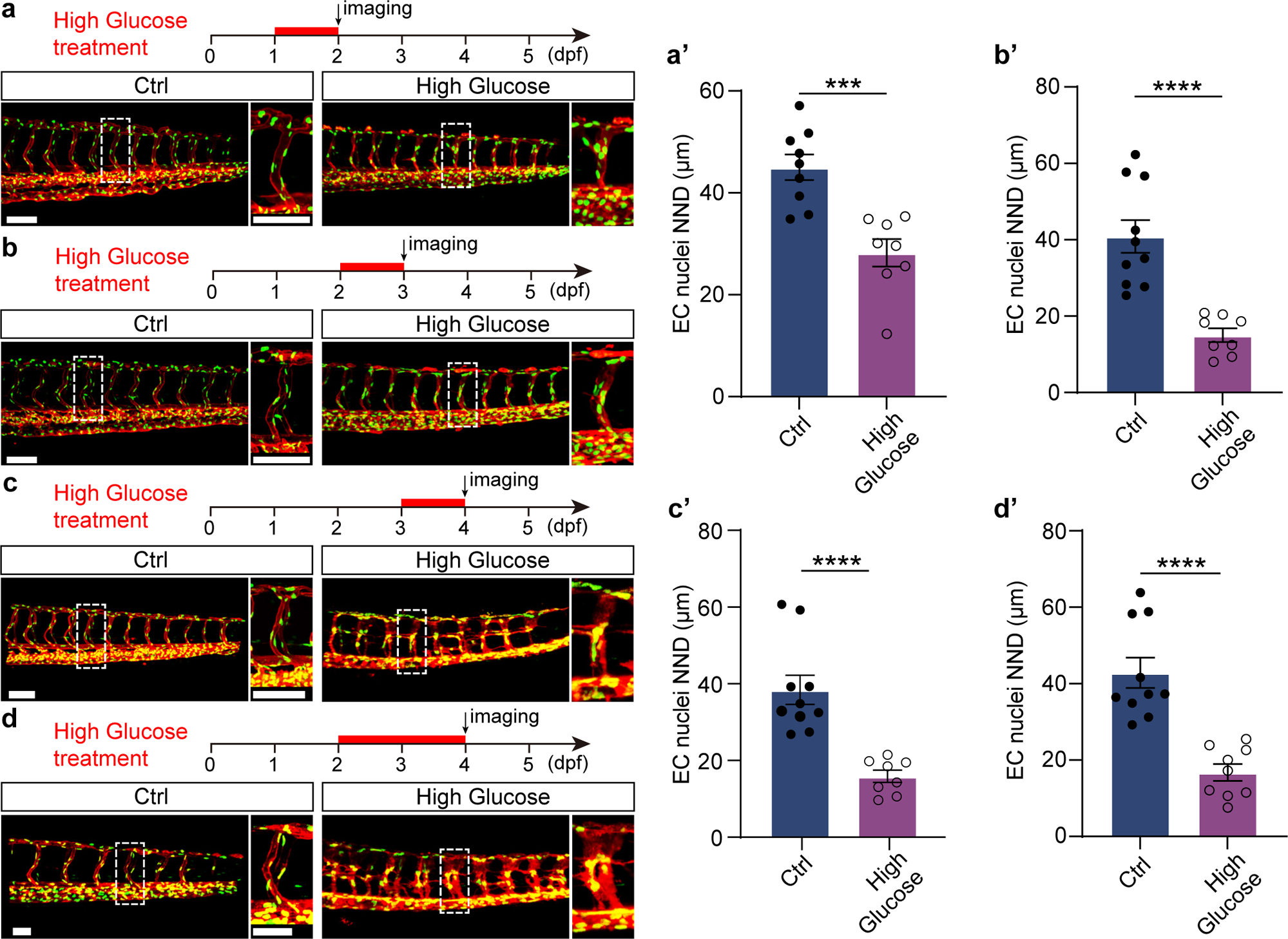
The aggregation of endothelial cell nuclei in zebrafish embryos was induced by high glucose shock treatment at different time windows. (a-d) Schematic diagrams of different time windows of high glucose shock treatment and confocal images. The right panels are the magnifications of the white dotted boxes. (a’-d’) Statistics of the ECs nuclei NND in different time windows in a-d, respectively. *t*-test. \*\*\**p*<0.001, \*\*\*\**p*<0.0001. Scale bars, 100 µm.

### High glucose treatment causes excessive migration of vascular endothelial nuclei in zebrafish embryos

To further investigate the cellular effects of high glucose treatment on the endothelial cells of zebrafish embryos, we performed time-lapse imaging to surveil the behavior of ECs nuclei from 63 to 72 hpf (Figure 3a-n). Compared to the control group, the EC nuclei in embryos subjected to high glucose treatment exhibited hyperactivity. In contrast, the ECs nuclei in the control embryos remained relatively stationary (Figure 3a’-g’, o), while those in the treated embryos displayed excessive migration (Figure 3h’-n’, p). The distance of EC nuclei migration (DNM) in the treated embryos was significantly greater than that observed in the control embryos (Figure 3q). Furthermore, the analysis of the number of ECs nuclei within different migration distance ranges revealed that the migration distance of ECs nuclei in control embryos was predominantly within the range of 10∼15 μm. In contrast, in the treated embryos, it was primarily within the range of 30∼40 μm, approximately three times greater (Figure 3r). The results suggested that high glucose shock treatment led to excessive migration of EC nuclei in zebrafish embryos.

**Figure 3.**
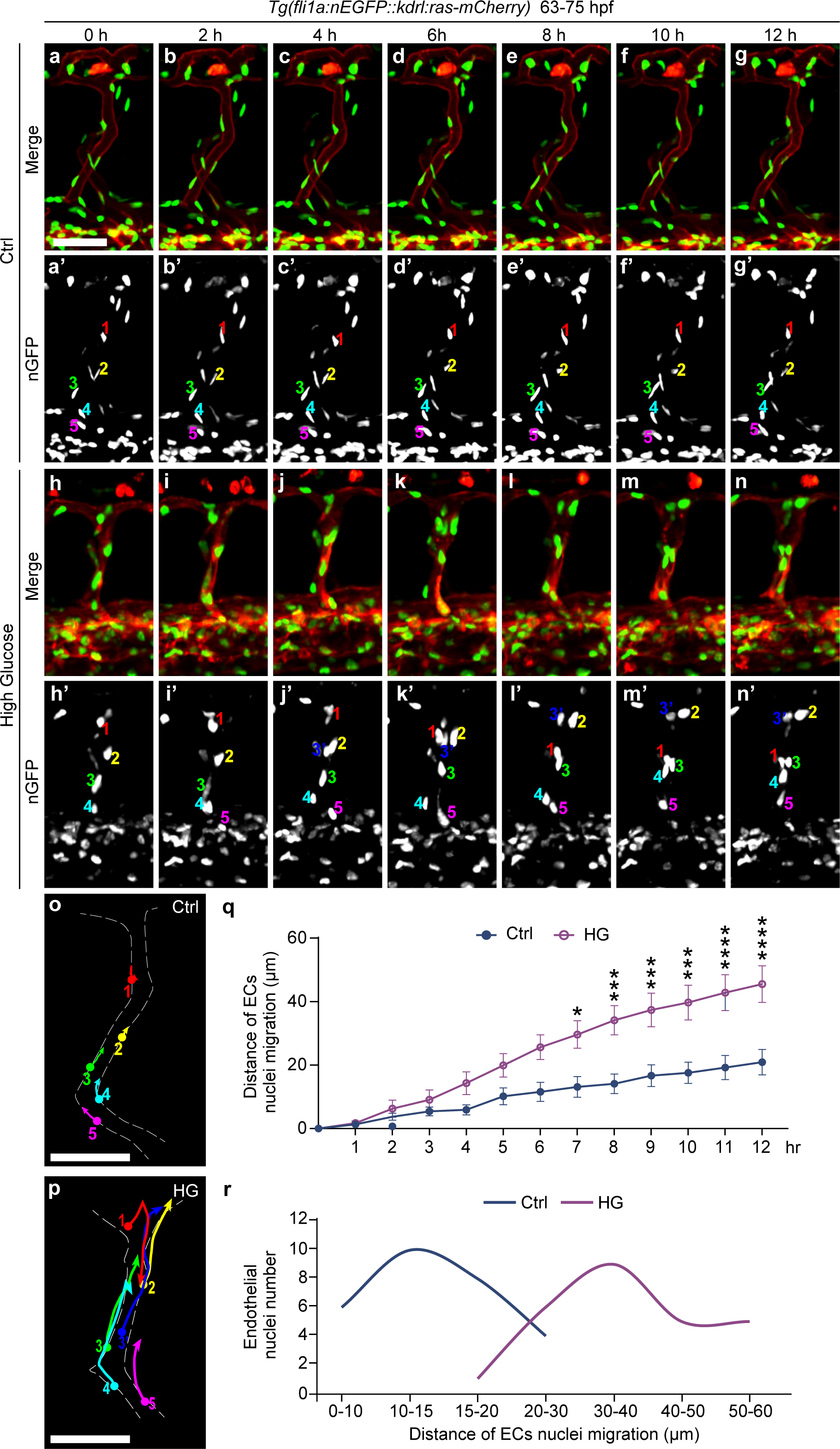
High glucose shock treatment causes excessive migration of vascular endothelial cell nuclei in ISVs of zebrafish embryos. (a-n) Still images from in vivo time-lapse imaging analysis of *Tg(fli1a:nEGFP::kdrl:ras-mCherry)* embryos from 63 to 75 hpf. (a’-n’) The nEGFP channel of the above panels, respectively. The numbers represent different endothelial cell nuclei in unilateral ISVs. (o-p) Diagrams of the migration routes of endothelial cell nuclei labeled in a’-n’. (q) Statistics of the distance of ECs nuclei migration (DNM) at different time stages in control embryos and high glucose shock treated embryos. Two-way ANOVA. (r) The number of endothelial nuclei in different migration distances of control embryos and high glucose-treated embryos. \**p*<0.05, \*\*\**p*< 0.001, \*\*\*\**p*< 0.0001. Scale bars, 50 µm.

### High glucose-induced aggregation of vascular endothelial nuclei via *foxo1a* downregulation in zebrafish embryos

Our previous work found that high glucose-induced excessive sprouting angiogenesis in zebrafish was mediated by down-regulation of *foxo1a*. It has been reported that FOXO1 was involved in the process of EC migration [29]. Hence, we hypothesis that high glucose shock induced the aggregation of vascular endothelial nuclei was also mediated by *foxo1a*. In order to verify our speculation, control zebrafish embryos were treated with AS1842856, a cell-permeable inhibitor that blocks the transcription activity of Foxo1 [30]. The results revealed that the inhibition of Foxo1 also resulted in the aggregation of EC nuclei in ISVs, resembling the phenotype observed in embryos subjected to high glucose shock. Moreover, the NND of EC nuclei in AS1842856 treated embryos was significantly lower compared to that of control embryos (Figure 4a-c). To further confirm the effect of *foxo1a*, we performed a rescue experiment by overexpressing *foxo1a* in zebrafish embryos. The *Tg(hsp70l:foxo1a*-6×His*-P2A-mCherry*) constructs and Tol2 transposase mRNA were co-microinjected into one-cell-stage embryos and followed by heat shock treatment at 24 hpf to overexpress Foxo1a. Next, the control and embryos overexpressing Foxo1a were treated with high glucose from 48 to 72 hpf. The results showed that overexpression of Foxo1a reduced the aggregation of vascular endothelial nuclei induced by high glucose (Figure 4d-e). Subsequently, we performed Foxo1 inhibition and wound healing assay in HUVECs. The cells were treated with 0.5 µM AS1842856 for 24 hours, and the wound healing assay result showed that Foxo1 inhibition promoted HUVECs migration compared to DMSO treatment (Figure 4f-h). These results suggest that high glucose may induce the nuclei aggregation and migration of ECs in zebrafish embryos through *foxo1a*.

**Figure 4.**
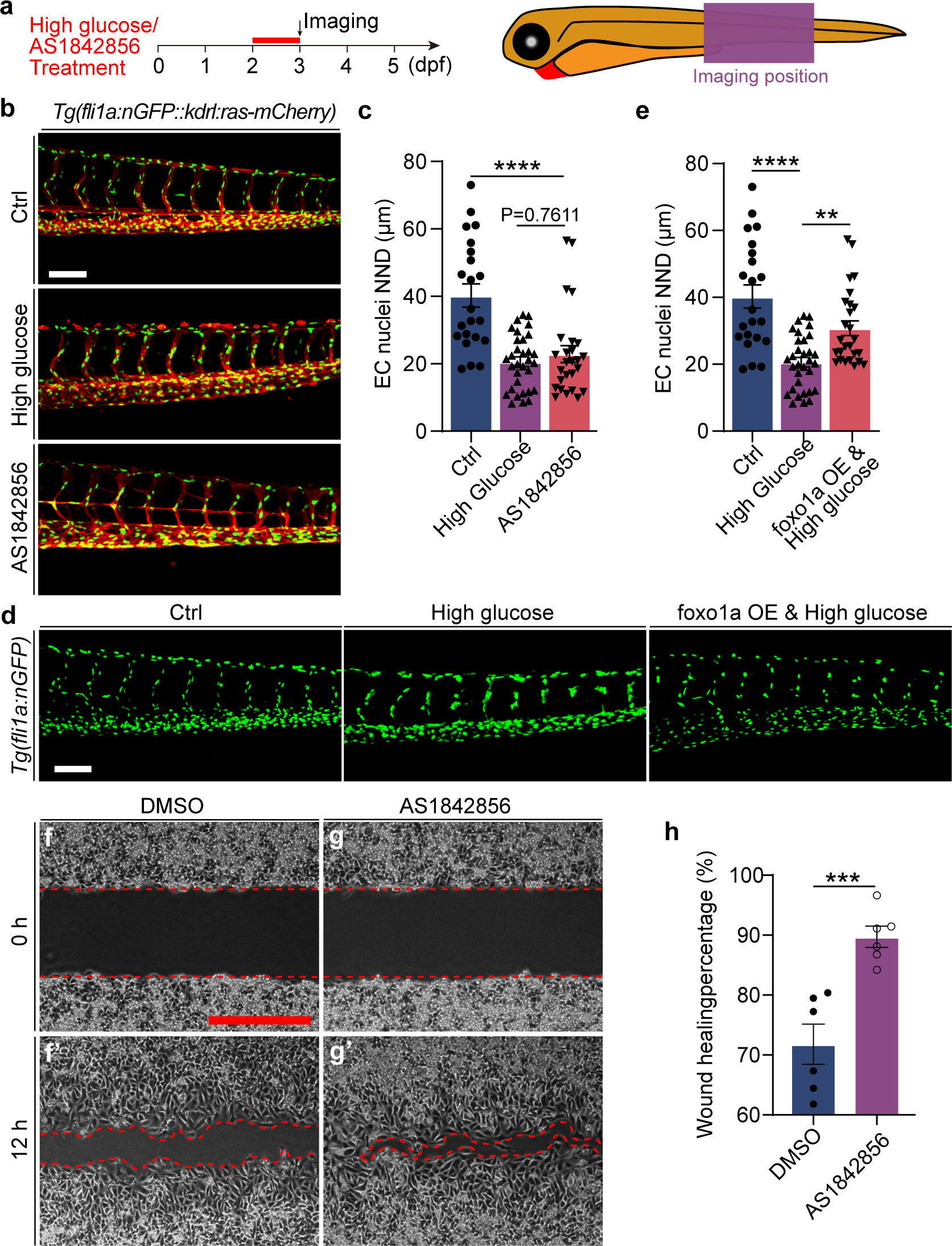
Foxo1 deficiency promotes HUVEC migration and results in the aggregation of endothelial cell nuclei in zebrafish embryos. (a) Schematic diagram of drugs treatment time window and confocal imaging region. (b) Confocal imaging analysis of control, high glucose shock, and AS1842856 treated *Tg(fli1a:nEGFP::kdrl:ras-mCherry)* embryos at 72 hpf. (c) Statistics of the ECs nuclei NND in control, high glucose shock, and AS1842856 treated embryos. *t*-test. (d) Confocal imaging analysis of ECs nuclei in ISVs in the control embryos, high glucose shock treated embryos, and *Tg(hsp70l:foxo1a*-6×His*-P2A-mCherry*) & high glucose shock treated embryos at 72 hpf. (e) Statistical analysis of the ECs nuclei NND in the control embryos, high glucose shock treated embryos, and *Tg(hsp70l:foxo1a*-6×His*-P2A-mCherry*) & high glucose shock treated embryos at 72 hpf. one-way ANOVA. (f-g’) The results of the wound healing assay showed that Foxo1 deficiency promotes HUVECs migration, in comparison with DMSO treated group. (h) Statistical analysis of the wound healing percentage in the DMSO group and AS1842856 treated group. *t*-test. \**p*<0.05, \*\*\**p*<0.001, \*\*\*\**p*<0.0001. Scale bars, 100 µm.

### Klf2a was significantly upregulated in arterial and capillary ECs

To identify the potential downstream factors for nuclei aggregation and migration of ECs in the embryos treated with glucose, we analyzed and compared the differentially expressed genes (DEGs) in arterial and capillary ECs of control and glucose-treated ECs, which was described in our previous work [28]. We performed GO analysis of significantly up-regulated genes, and the results revealed that these genes were enriched in several biological processes, including cellular catabolic process, intracellular transport, establishment of localization in cells, actin filament severing, etc. (Figure 5a). Subsequently, the up-regulated genes were further intersected with FOXO1 target genes and EC migration-associated genes, and we found *klf2a* (Figure 5b). ScRNA-seq data revealed that *klf2a* was significantly upregulated in arterial and capillary ECs after high glucose treatment (Figure 5c-c’). The *in-situ* hybridization experiment further confirmed the expression increase of *klf2a* following high glucose treatment (Figure 5d).

**Figure 5.**
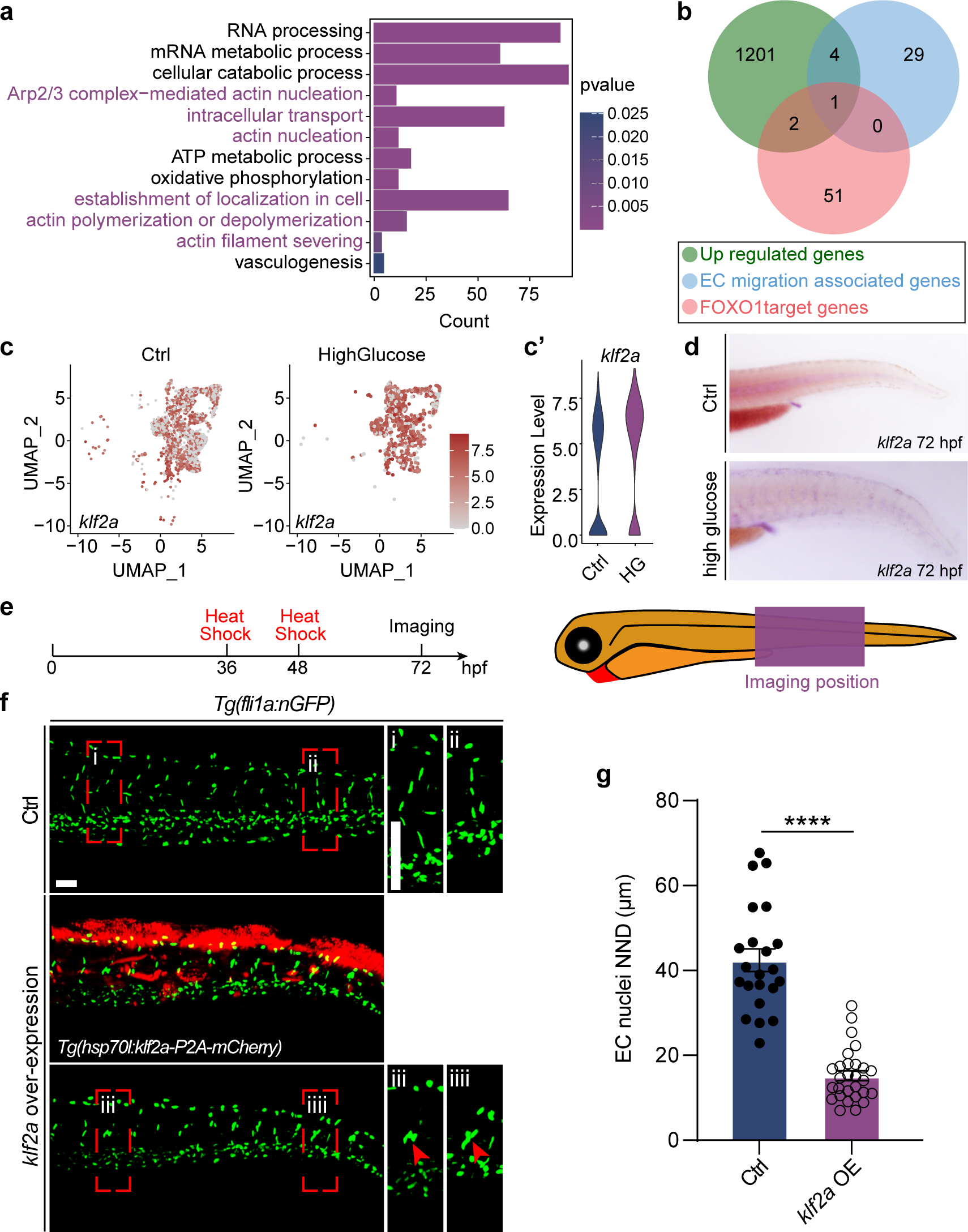
Klf2a was involved in the nuclei aggregation induced by high glucose shock treatment. (a) GO analysis of 1201 up-regulated genes in arterial and capillary ECs. (b) Venn diagram showing the overlap between FOXO1 target genes, EC migration-associated genes, and significantly up-regulated genes. (c) The feature plot of *klf2a* of control and high glucose group in arterial and capillary ECs. (c’) The violin plot of *klf2a* of control and high glucose group in arterial and capillary ECs. (d) Whole-mount in situ hybridization analysis of *klf2a* in control and high glucose treated embryos. (e) Schematic diagram of heat shock treatment and confocal imaging region. (f) Confocal imaging analysis of control embryos and *Tg(hsp70l:klf2a-P2A-mCherry*) embryos at 72 hpf. The right panels are the magnifications of the red dotted boxes. Arrowheads indicate the ECs nuclei. (g) Statistics of the ECs nuclei NND in control and *Tg(hsp70l:klf2a-P2A-mCherry*) embryos. *t*-test. \*\*\*\**p*<0.0001. Scale bars, 50 µm.

To verify whether the up-regulation of *klf2a* led to the nuclei aggregation and migration of ECs in zebrafish embryos, we performed gain-of-function experiments targeting *klf2a*. we conducted an experiment with overexpression of *klf2a*. We microinjected the Tg(hsp70l:klf2a-P2A-mCherry) constructs and Tol2 transposase mRNA into one-cell-stage embryos, followed by heat shock treatment at 36 and 48 hpf (Figure 5e). The findings revealed that *klf2a* overexpression also results in nuclei aggregation and aberrant migration of ECs (Figure 5f-g, movie 3), suggesting that the up-regulation of klf2a contributes to the observed phenotype in high glucose-treated embryos.

### High glucose-induced nuclei aggregation and migration through the *foxo1a-klf2a* pathway

To examine whether Klf2 is a downstream effector of Foxo1, we performed a series of experiments. we first investigated the impact of Foxo1 inhibition on Klf2 expression in zebrafish embryos and HUVECs. As expected, qPCR analysis revealed that inhibition of Foxo1 by AS1842856 resulted in up-regulation of KLF2 and klf2a expression in HUVECs and zebrafish embryos, respectively (Figure 6a-b). Next, we searched the FOXO1 target sequence through the JASPAR database (https://jaspar.genereg.net/) and identified three potential binding sites in the *klf2a* promoter region (Figure 6c-d). ChIP-PCR results showed that within 2 kb upstream of klf2a transcription start sites (TSS), a sequence of 5’-ATGTAAACATT-3’ at −1185 to −1195 nucleotides is a potential Foxo1a binding site (BS) of zebrafish, suggesting that Foxo1a might bind to the promoters of *klf2a* and regulate its transcription (Figure 6e).

**Figure 6.**
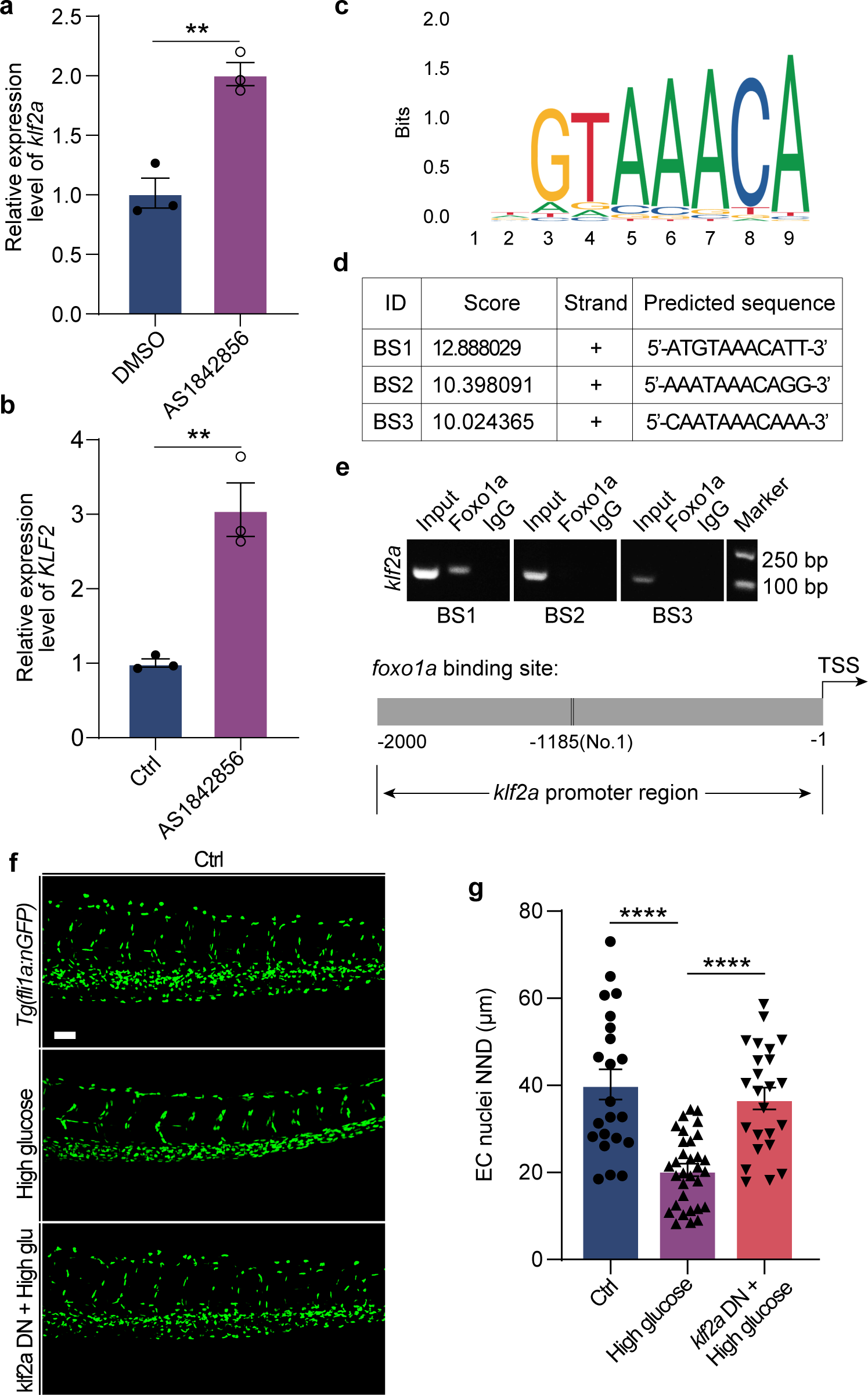
High glucose treatment induced nuclei aggregation through *foxo1a-klf2a* signal in zebrafish embryos. (a) Real-time PCR analysis of *klf2a* expression in control and AS1842856 treated embryos. t-test. (b) Real-time PCR analysis of KLF2 expression in HUVECs treated with DMSO and AS1842856. t-test. (c) A potential Foxo1 binding sequence presented in JASPAR database. (d) Three Foxo1a different candidate binding sites at the upstream of TSS of *klf2a* in zebrafish. (e) Results of the ChIP-PCR assay demonstrated that the predicted sequence “ATGTAAACATT” is a Foxo1a-binding site of *klf2a* in zebrafish. Schematic diagram showing the Foxo1a binding site in the *klf2a* promoter region. Input sonicated genomic DNA samples without immunoprecipitation as a positive control. IgG, sonicated, and IgG-immunoprecipitated genomic DNA samples as a negative control. TSS, transcription start site. (f) Confocal imaging analysis of control embryos, high glucose shock treated embryos, and *Tg(hsp70l:mApple-klf2a-DN)* & high glucose shock treated embryos at 72 hpf. (g) Statistical analysis of the ECs nuclei NND in the control embryos, high glucose shock treated embryos, and *Tg(hsp70l:mApple-klf2a-DN)* & high glucose shock treated embryos at 72 hpf. one-way ANOVA. \*\*\*\**p*<0.0001. Scale bars, 50 µm.

Additionally, we generated the *klf2a* dominant negative construct *Tg(hsp70l:mApple-klf2a-DN)* (Supplementary Figure 2) and microinjected the construct and Tol2 mRNA into the embryos at one-cell-stage, followed by heat shock treatment at 36 and 48 hpf. Subsequently, the embryos were treated with high glucose, and their phenotypes were examined using confocal imaging analysis. The results demonstrated that Klf2a deficiency significantly restored the ECs nuclei distribution (Figure 6f-g). Collectively, these results suggest that high glucose shock treatment might induce the aggregation of ECs nuclei in zebrafish embryos via the *foxo1a-klf2a* pathway.

## Discussion

Microvascular complications in diabetes are those long-term damages that impact small blood vessels. The etiology of this condition involves a confluence of decreased eNOS, oxidative stress, and the generation of advanced glycosylation end products [31–33]. Diabetic retinopathy, a prevalent and illustrative microvascular complication, initially manifests through the existence of microaneurysms [34, 35]. Microaneurysms are tiny areas of swelling in the blood vessels. A microaneurysm consists of several crowded nuclei of endothelial cells. In this study, we observed an aggregation of vascular endothelial nuclei in the ISVs of zebrafish embryos induced by high glucose, which is similar to the crowded nuclei of endothelial cells in microaneurysms (Figure 7). Further observation revealed hyperaction and excessive migration of the endothelial cells. Consequently, we deduce that the aggregation of the endothelial nuclei may result from the aberrant migration of these cells. It is recognized that both proliferation and migration are crucial for the process of angiogenesis. However, excessive proliferation and migration can be detrimental and contribute to a range of malignant ailments, such as various diabetic vascular complications, atherosclerosis, restenosis, and cancers [36, 37]. Here, our findings offer new evidence elucidating the impact of high glucose levels on the vascular status via regulating the behaviors of the endothelial cells.

**Figure 7.**
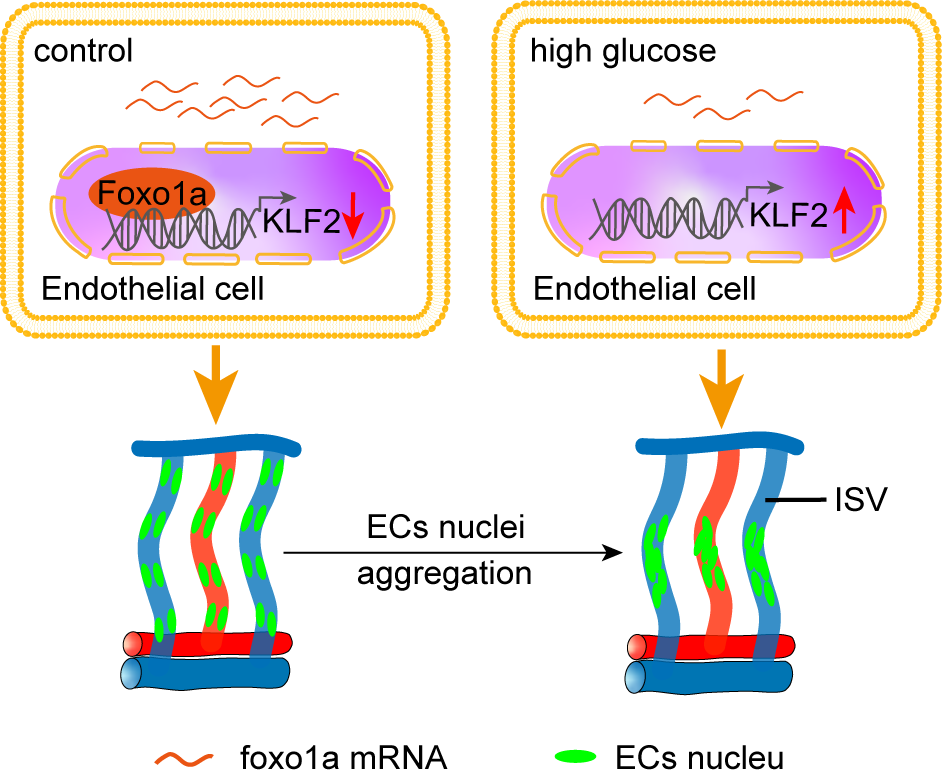
Working model of high glucose treatment induced ECs nuclei aggregation in ISVs of zebrafish embryos.

Forkhead box protein O1 (FOXO1), also known as forkhead in rhabdomyosarcoma (FKHR), is a member of the forkhead transcription factor family. FOXO1 plays important roles in the regulation of cell proliferation, metabolism, and stress resistance [38–40]. FOXO1 is primarily regulated by the insulin/PI3K/Akt signaling pathway. Activation of PI3K/AKT leads to the dephosphorylation of FOXO1, results in its translocation from the cytoplasm to the nucleus, and initiates transcription of target genes [41–43]. FOXO1 is involved in the regulation of EC metabolism, sprouting, proliferation, and migration. Sustained FOXO1 activation causes a thin and hyperpruned vascular network with fewer ECs [44–46]. Endothelial-specific FOXO1 deficiency promotes vascular growth in the adipose tissue of obese mice [47]. Accumulating evidence has demonstrated that FOXO1 is also involved in the development and progression of diabetes. FOXO1 activation impairs eNOS function in vascular endothelial cells resulting in endothelial dysfunction [48]. FOXO1 inhibition promotes angiogenesis and recovery of EC function in diabetic mice [49]. FOXO1 also plays an important role in the pathogenesis of diabetic cardiomyopathy [50]. There have also been some studies showing the role of FOXO1 in microvascular complications [51–53]. Previous research has documented the significant involvement of Foxo1 in the regulation of gluconeogenesis and glycogenolysis through the insulin signaling pathway [54, 55]. Activation of Foxo1 due to insulin resistance has been observed to exacerbate diabetic cardiomyopathy, whereas its deficiency has been shown to effectively mitigate heart failure and restore cardiac function in db/db and high-fat diet mice [56]. In zebrafish, *foxo1* has two duplicate genes named *foxo1a* and *foxo1b*. Although the role of *foxo1a* and *foxo1b* in zebrafish vasculature has been little studied, a recent study has reported that *foxo1* is involved in embryonic vascular development [57]. Investigations have revealed that the depletion of foxo1, specifically in endothelial cells, enhances metabolic activity and promotes angiogenesis in both normal and obese mice, thus indicating its inhibitory role in angiogenesis, which aligns with our own findings. In this study, our investigation employed techniques such as single-cell sequencing analysis and ChIP-PCR to provide evidence suggesting that Foxo1a may play a role in regulating endothelial cells’ nuclei aggregation in the ISVs of zebrafish embryos treated with high glucose. This regulation is achieved through the inhibition of *klf2a* expression. Notably, KLF2 is a crucial transcriptional factor in endothelial cells that controls nitric oxide production, a molecule with significant implications in renal disease [58]. The dysregulation of KLF2 has been shown to be responsible for cardiac microvascular disease in diabetes, specifically linked to defects in monocyte adhesion and migration in endothelial cells [59]. Previous research has identified the reciprocal regulation of Klf2 by Foxo1 in HUVECs, suggesting a potential mechanism for diabetic endothelial dysfunction [60]. In our study, we utilized ChIP-PCR and dominant negative experiments to confirm the involvement of *klf2a* and its interaction with Foxo1 in endothelial cells of zebrafish embryos exposed to high glucose treatment.

To date, existing literature has indicated a correlation between endothelial dysfunction in microvascular complications and heightened oxidative stress, reduced release of nitric oxide, activation of protein kinase C, and heightened production of inflammatory factors [61]. However, understanding the underlying cellular and molecular mechanisms responsible for diabetic microvascular complications remains limited. This study, for the first time, demonstrates that exposure to high glucose treatment induces aggregation of endothelial cell nuclei and aberrant migration of endothelial cells *in vivo*. Furthermore, it has been determined that this particular phenotype induced by high glucose is mediated through the *foxo1a-klf2a* axis. In summary, our findings contribute a novel insight into the mechanism of microvascular complications in hyperglycemia.

## Materials and Methods

### Zebrafish

*Tg(kdrl:ras-mCherry)* and *Tg(fli1a:nEGFP)* transgenic zebrafish were used in this study, in which endothelial cell membrane was labeled with mCherry and endothelial nucleus was labeled with EGFP. Zebrafish embryos for Whole-mount in situ hybridization were obtained through natural mating (AB line). Zebrafish embryos were treated with 0.2 mM 1-phenyl 2-thiourea (Sigma-Aldrich, P7629) after 24 hpf to prevent pigment formation. According to our previous work, all of these embryos and adult fish were maintained under standard conditions [62].

### Drug treatment

Zebrafish embryos with developmental defects or delays were removed at 8 hpf and then raised in 12-well plates at 10 embryos per well at 28.5°C. Embryos were treated with 300 mM D-glucose (Sigma, G7021) at different time windows. AS1842856 (MCE, HY-100596) powder was dissolved in DMSO into a 10 mM stock solution, stored at 80°C, and used at 1 µM for zebrafish embryos and 0.1 µM for HUVEC. The same concentration of DMSO was used as a negative control.

### Glucose concentration measurement

Glucose concentration in the embryos was measured as described in our previous work [28]. Briefly, embryos were selected and transferred to 24-well plates (ten embryos per well) and immersed in high glucose solution at 48 hpf. For glucose concentration measurement, embryos (n=20) were transferred to a new 1.5 mL tube, rinsed three times with 1×PBS, and then discarded the PBS as much as possible. Subsequently, embryos were homogenized using a hand homogenizer and centrifuged at 14,000×g for 10 min. 1.5 μL of the supernatant was used to measure the total free-glucose level using a glucometer (Baye, 7600P).

### Whole-mount in situ hybridization (WISH)

The whole mount in situ hybridization and the preparation of an antisense RNA probe were performed as described in the previous protocol [63]. The *klf2a* cDNA fragment was cloned with the specific primers listed below using the wild-type embryo (AB) cDNA library. The antisense probe was synthesized using the in vitro DIG-RNA labeling transcription Kit (Roche, 11175025910) with linearized pGEM-T easy vector containing *klf2a* fragment as the templates. The synthesized probe was purified with LiCl (Invitrogen, AM9480) and diluted to 1 ng/µL for hybridization. Zebrafish embryos at 72 hpf were collected and fixed with 4% paraformaldehyde (PFA) overnight at 4°C and then dehydrated with gradients of methanol and stored at −20°C in 100% methanol for subsequent analysis. The hybridization result was detected with anti-DIG-AP antibody (1:2000, Roche, 11093274910) and NBT/BCIP (1:500, Roche, 11681451001). After hybridization, images of the embryos were captured with an Olympus stereomicroscope MVX10.

*klf2a*-probe-forward:5’-GAA TGA GGA CTA CCC GGA CC-3’

*klf2a*-probe-reverse:5’-CCT CAT TCT GCA CTC GGA TGA-3’

### RNA isolation, reverse transcription, and quantitative RT-PCR (qRT-PCR)

Total RNA was isolated with TRIzol Reagent (Invitrogen, 15596026) and stored at −80°C. Afterward, the cDNA was synthesized using the HiScript III First Strand cDNA Synthesis Kit (Vazyme, R312-01) and stored at −20°C. Quantitative PCR was carried out using the Taq Pro Universal SYBR qPCR Master Mix (Vazyme, Q712-02) on the basis of the manufacturer’s instructions. Primers for Real-time PCR analysis are as follows:

KLF2-Homo-qPCR-forward:5’-CTA CAC CAA GAG TTC GCA TCT G-3’

KLF2-Homo-qPCR-reverse:5’-CCG TGT GCT TTC GGT AGT G-3’

*klf2a*-Dre-qPCR-F: 5’-CTC TAG AGC TGG ACG CCA AA-3’

*klf2a*-Dre-qPCR-R: 5’-GAT AGG GCT TCT CGC CTG TG-3’

### Plasmid construction and microinjection

The coding sequence of *klf2a* and the sequence of *foxo1a*-6×His were synthesized and inserted respectively into *hsp70l:MCS-P2A-mCherry* vector by Genewiz Co., Ltd. (Suzhou, China). The sequence of *klf2a*-DN was designed referring to the previous work[64] and added to the C-terminal of mApple fluorescent protein through gene synthesis to get the construction of *Tg(hsp70l:mApple-klf2a-DN)*. For microinjection, 60 ng Tol2 transposase and 75 ng plasmid were mixed well in 5 µL water, respectively, and a 2 nL mixture was microinjected into zebrafish embryos at the one-cell stage. For heat shock treatment, embryos were transferred to a 1.5 ml centrifuge tube, incubated for 1 hour at 37°C, and then raised in the dish with new egg water at 28.5°C.

### ChIP-PCR

Embryos injected with *Tg(hsp70l:foxo1a-6×His-P2A-mCherry)* were collected at 72 hpf after heat shock treatment. The ChIP-PCR assay was performed using the Chromatin Immunoprecipitation (ChIP) Assay Kit (Millipore, 3753379) according to the manufacturer’s instructions. The genomic DNA crossed with Foxo1a protein was immunoprecipitated by Ni NTA Beads (Smart-Lifesciences, SA004100) following the manufacturer’s instructions. Antibody against lgG was used as a negative control. The semiquantitative PCR was performed with KODfx (TOYOBO, KFX-101) at the following conditions: 94°C for 5 min; 35 cycles of 98°C for 10 s, 55°C for 30 s, 68°C for 10 s; 68°C for 10 min. The PCR primers used for the predicted binding sites (BS) are as follows:

*klf2a*-BS1-forward:5’-CTG ACT TCA ACT GTA TAT AAA GGC-3’

*klf2a*-BS1-reverse:5’-GTT GAG ATA AAA TCT TCA TAA TGC-3’

*klf2a*-BS2-forward:5’-ATA ATC TCT CCG TTA AAC GGG-3’

*klf2a*-BS2-reverse:5’-GGT TAT GTG CCT ATG TGT ACA GTG-3’

*klf2a*-BS3-forward:5’-CCA CTA TAC CTA AGC TAT TAT GTC-3’

*klf2a*-BS3-reverse:5’-TGA TCA TGC ATT TCG ATG GTT-3’

### Cell culture and wound-healing assay

HUVECs were cultured according to the protocols described previously[65, 66]. For the wound-healing assay, a dish with a defined cell-free gap (Ibidi, 80206) was used to create the gap following the manufacturer’s instructions. HUVECs were treated with 1 μg/ml Mitomycin C (Sigma-Aldrich, M5353) for 1 hour before the experiment to inhibit the effects of cell proliferation. Images were captured at 0 h, 6 h, 12 h, and 18 h after drug treatment with an Olympus microscope IX73, and the scratch area was measured with ImageJ software.

### Confocal imaging and quantitative analysis

For confocal imaging, zebrafish embryos were anesthetized with egg water/0.16 mg/ml tricaine (Sigma-Aldrich, A5040) and embedded in 0.7% low melting agarose. Living imaging was performed with Nikon A1R confocal microscopy. For time-lapse imaging, embryos were placed on a platform at a constant temperature of 28.5°C. All of the data was measured with ImageJ software. For time-lapse data, using the dorsal aorta as a reference, and then the distance of the endothelial nucleus migration was measured by the shift of the nucleus in the before and after images. The range (R) and the standard deviation (SD) of the distance from the nucleus to the midline (DNTM) per ISV were calculated using the following formula:

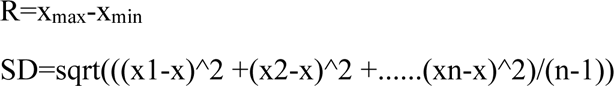

### Statistical analysis

Statistical analysis was performed with student’s t-test or One-Way ANOVA. All data is presented as Mean ± SEM, and *p* < 0.05 was considered to be statistically significant.

## Availability of data and material (data transparency)

All the experimental materials generated in this study are available from the corresponding authors upon reasonable request.

## Authors’ contributions

Dong Liu and Liping Chen conceived and designed the experiments and wrote the manuscript. Xiaoning wang, Xinyi Kang, and Bowen Li performed the experiments and analyzed the data. All authors read and approved the final manuscript.

## Ethics approval

All zebrafish experimentation was carried out following the NIH Guidelines for the care and use of laboratory animals (http://oacu.od.nih.gov/regs/index.htm) and ethically approved by the Administration Committee of Experimental Animals, Jiangsu Province, China (Approval ID: 20180905-002).

---

**Supplementary Figure 1.**
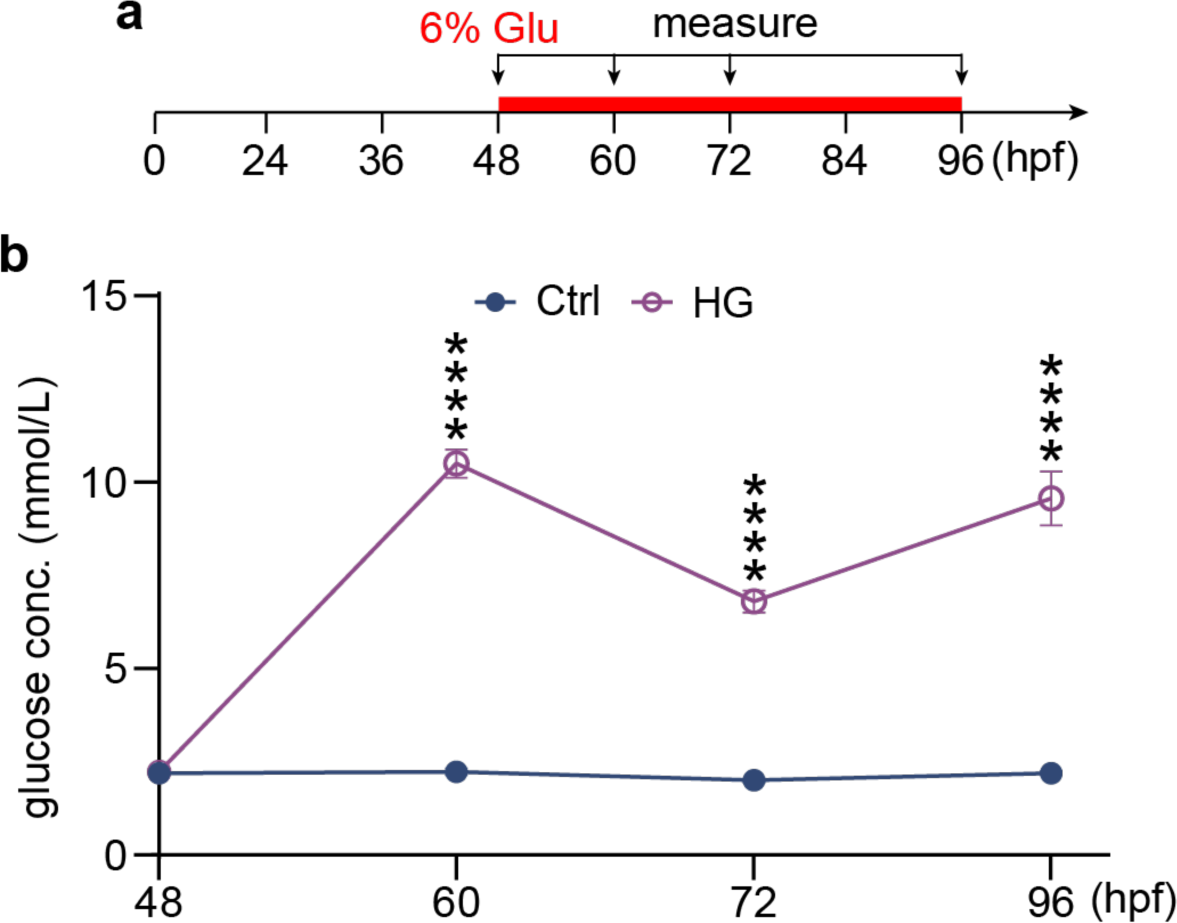
Total glucose concentrations at different development stages in control and high glucose-treated embryos. (a) A diagram showing the glucose treatment time window and concentration measuring time point. (b) Statistical analysis of the glucose concentration in control and high glucose-treated embryos. one-way ANOVA, ****p<0.0001.

**Supplementary Figure 2.**
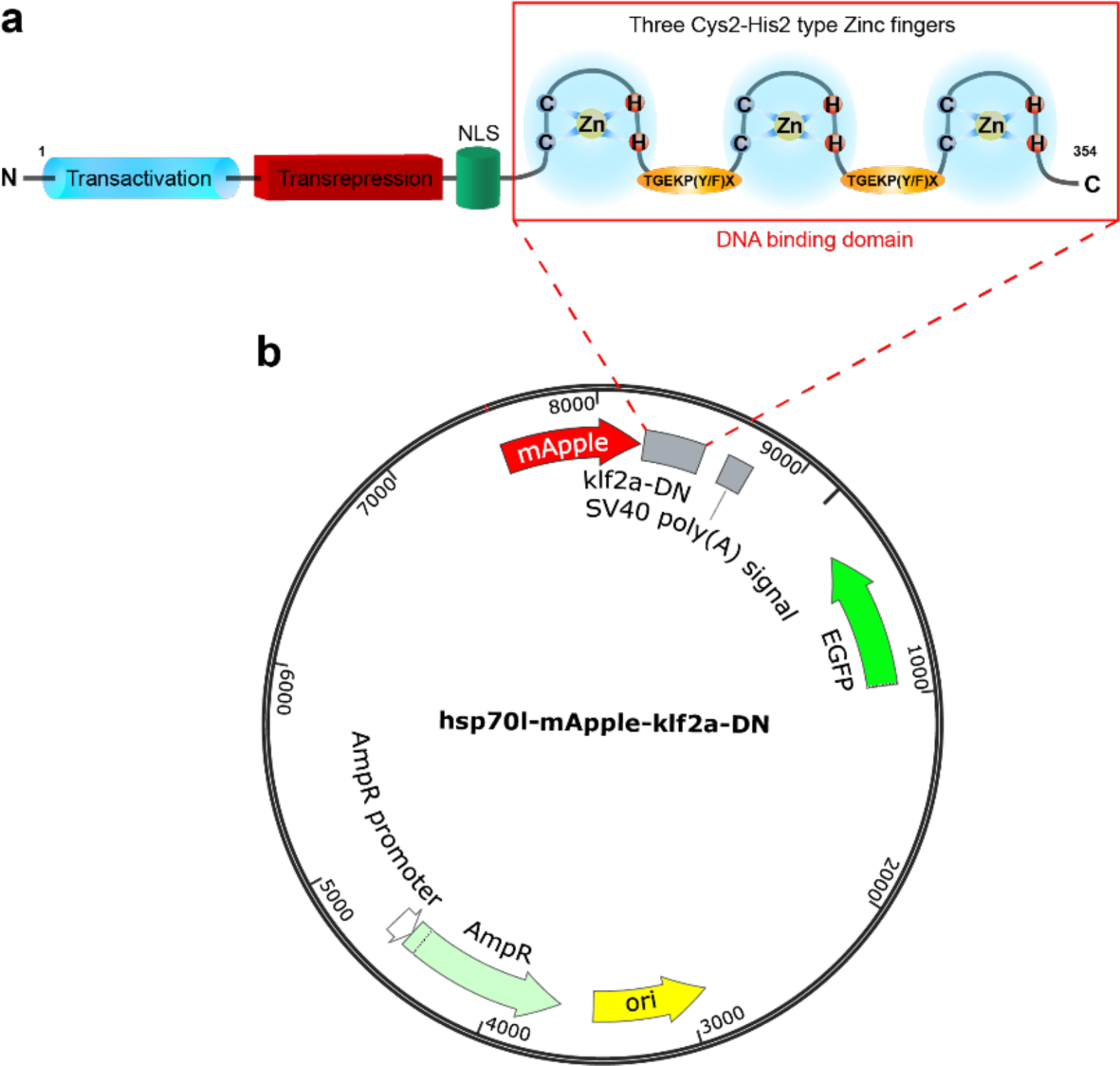
Generation of the *klf2a* dominant-negative construct. (a) Schematic representation of the klf2a structure. (b) Structural map of hsp70l:mApple-klf2a-DN construct.

**Video 1.** In vivo time-lapse imaging of control *Tg(fli1a:nEGFP)* embryos from 63 to 75 hpf.

**Video 2.** In vivo time-lapse imaging of high glucose treated *Tg(fli1a:nEGFP)* embryos from 63 to 75 hpf.

**Video 3.** In vivo time-lapse imaging of *klf2a* over expression *Tg(fli1a:nEGFP)* embryos from 63 to 75 hpf.

